# A Panel of Human Monoclonal Antibodies for Tracking the Antigenic Evolution of Influenza H5N1 Clade 2.3.4.4b

**DOI:** 10.64898/2026.05.14.725215

**Authors:** Hsiang Hong, Nicholas C. Morano, Madeline Wu, Sue Chong, Jian Yu, Yaoxing Huang, Manoj S. Nair, Zhiteng Li, Chih-Chen Tzang, Jordan E. Becker, Chi Wang, Lawrence Shapiro, Peter D. Kwong, Yicheng Guo, David D. Ho

**Affiliations:** Aaron Diamond AIDS Research Center, Columbia University Vagelos College of Physicians and Surgeons, New York, NY 10032, USA; Department of Microbiology, National Taiwan University College of Medicine, Taipei, Taiwan; Zuckerman Mind Brain Behavior Institute, Columbia University, New York, NY 10027, USA; Department of Biochemistry and Molecular Biophysics, Columbia University, New York, NY 10027, USA; Division of Infectious Diseases, Department of Medicine, Columbia University Vagelos College of Physicians and Surgeons, New York, NY 10032, USA; Department of Microbiology and Immunology, Columbia University Vagelos College of Physicians and Surgeons, New York, NY 10032, USA; Wu Center for Pandemic Research, Columbia University Vagelos College of Physicians and Surgeons, New York, NY 10032, USA, a unit of the Pandemic Research Alliance

**Keywords:** antigenic evolution, clade 2.3.4.4b, H5N1, influenza virus, monoclonal antibodies, hemagglutinin

## Abstract

The ongoing panzootic of clade 2.3.4.4b H5N1 influenza has driven widespread transmission among birds, mammals, and livestock, underscoring the need for tools to interpret its rapid antigenic evolution. Here, we generated and characterized 19 human monoclonal antibodies that potently neutralize clade 2.3.4.4b viruses and collectively recognize five major, partially overlapping antigenic regions of the H5 hemagglutinin. Neutralization profiling and structural analyses revealed distinct patterns of antibody breadth and escape, identifying both clade 2.3.4.4b restricted and broadly neutralizing antibodies targeting conserved surfaces across divergent H5N1 lineages. By incorporating these antibodies into an expanded 32-mAb reference panel, we identified antibody-defined antigenic differences among recent human isolates between cattle-associated B3.13 and poultry-associated D1.1 genotypes. Together, this study establishes a structurally informed, epitope-resolved H5 hemagglutinin antibody reference panel for interpreting emerging HA mutations and monitoring antigenic changes in circulating H5 viruses.

## Introduction

Highly pathogenic avian influenza H5N1 viruses pose a pandemic threat due to their high virulence, broad host range, and ongoing global circulation in avian and mammalian species^1,2^. Since the first human fatality during the 1997 Hong Kong outbreak^3,4^, H5N1 has caused sporadic zoonotic disease with a case fatality rate of ∼50% among confirmed cases^5^. For more than two decades, human infections remained rare and geographically constrained. However, the emergence of clade 2.3.4.4b in 2020 marked a profound shift in the epidemiology of H5N1 viruses. This clade has rapidly driven an unprecedented panzootic, establishing sustained transmission in wild birds, devastating poultry populations across multiple continents, and expanding into a broad range of mammalian hosts^6–8^. By the end of 2025, this wave of viral spread had also resulted in 72 laboratory-confirmed human infections in the United States^9^ and Canada^10^.

Among newly affected mammals, dairy cattle have emerged as a particularly consequential host for clade 2.3.4.4b H5N1. Genomic analyses indicate that genotype B3.13 was introduced into U.S. dairy cattle from wild birds and subsequently spread within and between herds across multiple states^11^. More recently, genotype D1.1, which has circulated extensively among poultry, was also introduced into dairy cattle through an independent spillover event^12,13^. In North America, human infections linked to B3.13-exposed cattle have been predominantly mild and often limited to conjunctivitis^5^, whereas D1.1 has been implicated in severe cases^10^ including three fatalities^14–16^. Although clinical outcomes are multifactorial and may be influenced by mutations in PB2 and other internal viral genes^17,18^, these observations underscore the diverse consequences of H5N1 spillover into humans. Independently of clinical disease severity, the continued diversification of circulating H5N1 viruses raises important questions regarding changes in their antigenic properties. Understanding how naturally occurring HA substitutions influence antibody recognition is therefore essential for characterizing the antigenic evolution of clade 2.3.4.4b viruses^19,20^.

Monoclonal antibodies (mAbs) are powerful tools for characterizing the antigenic properties of influenza hemagglutinin (HA) at the molecular level and connecting sequence changes to escape from immune pressure^21–23^. Several major vulnerable sites on H5 HA have previously been defined using mAbs derived from infections with earlier H5 viruses^24^, providing a foundational framework for understanding antibody recognition of H5 HA. More recently, potent mAbs against clade 2.3.4.4b viruses have been described following either historical H5N1 vaccination or immunization with a single clade 2.3.4.4b HA^25,26^. However, these antibody collections represent a limited range of epitope specificities, potentially restricting their utility for comprehensively characterizing antigenic differences among circulating clade 2.3.4.4b viruses. In parallel, although numerous potent antibodies against earlier H5 strains have been described^27–29^, many were isolated prior to the emergence of clade 2.3.4.4b and exhibit reduced neutralizing activity against recent isolates^30^. Hence, expanding the panel of antibodies recognizing diverse HA antigenic regions could facilitate systematic characterization of the ongoing antigenic evolution of clade 2.3.4.4b viruses.

In this study, we generated and characterized a panel of human monoclonal antibodies against the HA of the current panzootic clade 2.3.4.4b strain A/Texas/37/2024. By immunizing VelocImmune mice^31^ with four H5 antigens derived from WHO candidate vaccine viruses, including a clade 2.3.4.4b strain, we isolated 19 human mAbs spanning diverse germlines, neutralization potencies, and epitopes. Through binding and neutralization assays against representative avian and mammalian H5N1 isolates, competition ELISAs, and high-resolution cryo-electron microscopy (cryoEM), we characterized antibodies recognizing several major, partially overlapping antigenic regions of the HA head, as well as the conserved stem. We subsequently incorporated these antibodies into an expanded 32-mAb reference panel to compare the antigenic profiles of representative B3.13 and D1.1 viruses from recent human infections, revealing genotype-associated patterns of monoclonal antibody escape.

## Results

### Clade 2.3.4.4b H5N1 escapes most previously isolated HA head-directed mAbs

To track the antigenic evolution of recent clade 2.3.4.4b H5N1 viruses, we first assembled a panel of 14 previously published mAbs, including 7 antibodies directed to the HA head, 4 targeting the HA2 stem region, and 3 recognizing the anchor epitope at the membrane-proximal base of HA (**Table S1**). Stem-directed mAbs largely retained neutralizing activity against both a historical clade 0 virus and a representative clade 2.3.4.4b virus, consistent with the targeting of highly conserved functional regions within the HA stem. In contrast, most head-directed mAbs bound regions that typically accumulate antigenic drift mutations and showed markedly reduced potency against clade 2.3.4.4b. Only antibody 65C6^28^ maintained potent neutralization of both clade 0 and clade 2.3.4.4b viruses, highlighting substantial antigenic differences between historical and recent H5 HA head epitopes and the scarcity of broadly reactive head-directed mAbs.

### Isolation of H5N1-neutralizing antibodies from immunized VelocImmune^®^ mice

To build a comprehensive panel of human mAbs against recent H5N1 strains, 15 VelocImmune mice (#1-#15) were first immunized with DNA plasmids encoding HAs from four representative clades, including 0, 7.1, 2.2.1, and 2.3.4.4b, selected from WHO candidate vaccine viruses to encompass the diversity and broad antigenic spectrum of H5 (**Figure 1a**). Mice received two DNA primes followed by three boosts with recombinant HA trimer over a 30-week schedule (**Figure S1a**). Plasmids were administered into four limb sites simultaneously (**Figure S1b**). Serum neutralization titers against matched pseudoviruses were measured two weeks after each immunization; by week 8, sera exhibited robust neutralizing activity against all four immunization strains (**Figure 1b**).

**Figure 1.**
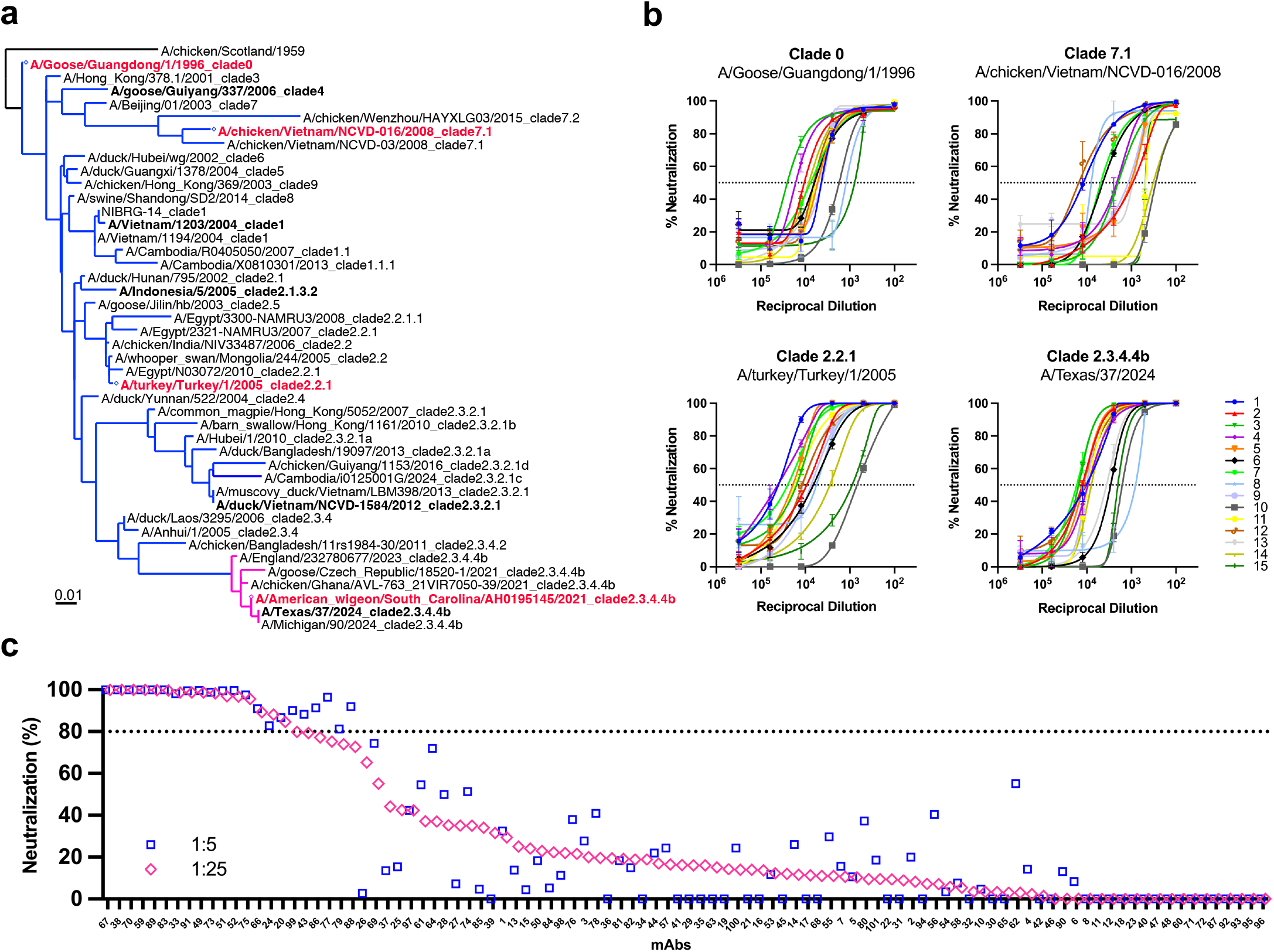
Isolation of H5N1-neutralizing monoclonal antibodies from VelocImmune® mice. a. Phylogenetic tree of WHO candidate vaccine viruses used in this study; immunization strains are highlighted in red, and viruses tested are shown in red or bold. b. Serum neutralization titers against immunization strains measured at week 8. c. Supernatant neutralization screening of selected monoclonal antibodies against A/Texas/37/2024.

Six mice (ID: #2, #3, #4, #5, #7, #9) with the highest serum neutralizing titers were selected for antigen-specific B cell retrieval. Splenocytes and peripheral blood mononuclear cells (PBMCs) were harvested and stained with fluorescently labeled HA probes from clade 0 (A/goose/Guangdong/1/1996) and clade 2.3.4.4b (A/Texas/37/2024). Antigen-specific B cells were sorted for 10x Genomics single-cell sequencing to obtain paired heavy- and light-chain variable regions, similar to what we did previously for isolating mAbs to SARS-CoV-2 spike^23,32–34^.

A total of 101 candidate mAbs were selected based on their clonal expansion, heavy-chain somatic hypermutation (SHM; >5%) and heavy-chain CDR3 length (>10 amino acids). These antibodies were cloned, expressed, and screened for binding and neutralization. While more than half bound to clade 2.3.4.4b HA (**Figure S1c**), 22 mAbs achieved ≥80% inhibition of A/Texas/37/2024 pseudovirus at a 1:25 dilution of culture supernatant (**Figure 1c**). Among these, the top 19 mAbs exhibiting the strongest pseudovirus neutralization activity were selected for downstream characterization of antibody expression, HA binding, and virus neutralization.

### Neutralization potency and breadth of mAbs

To evaluate the neutralizing activity of this panel, all 19 mAbs (hereafter referred to as the H- series antibodies) were first tested against the circulating clade 2.3.4.4b virus A/Texas/37/2024. All potently neutralized this strain in both pseudovirus (PV) and authentic live virus (LV) assays, with IC50 values ranging from ∼0.5 ng/mL to ∼1 µg/ml (**Figure 2a**). The IC50 values from the two assays were strongly correlated (r = 0.90), indicating that the PV assay could serve as a reliable surrogate for inhibitory activity against authentic H5N1, albeit at lower IC50 values.

**Figure 2.**
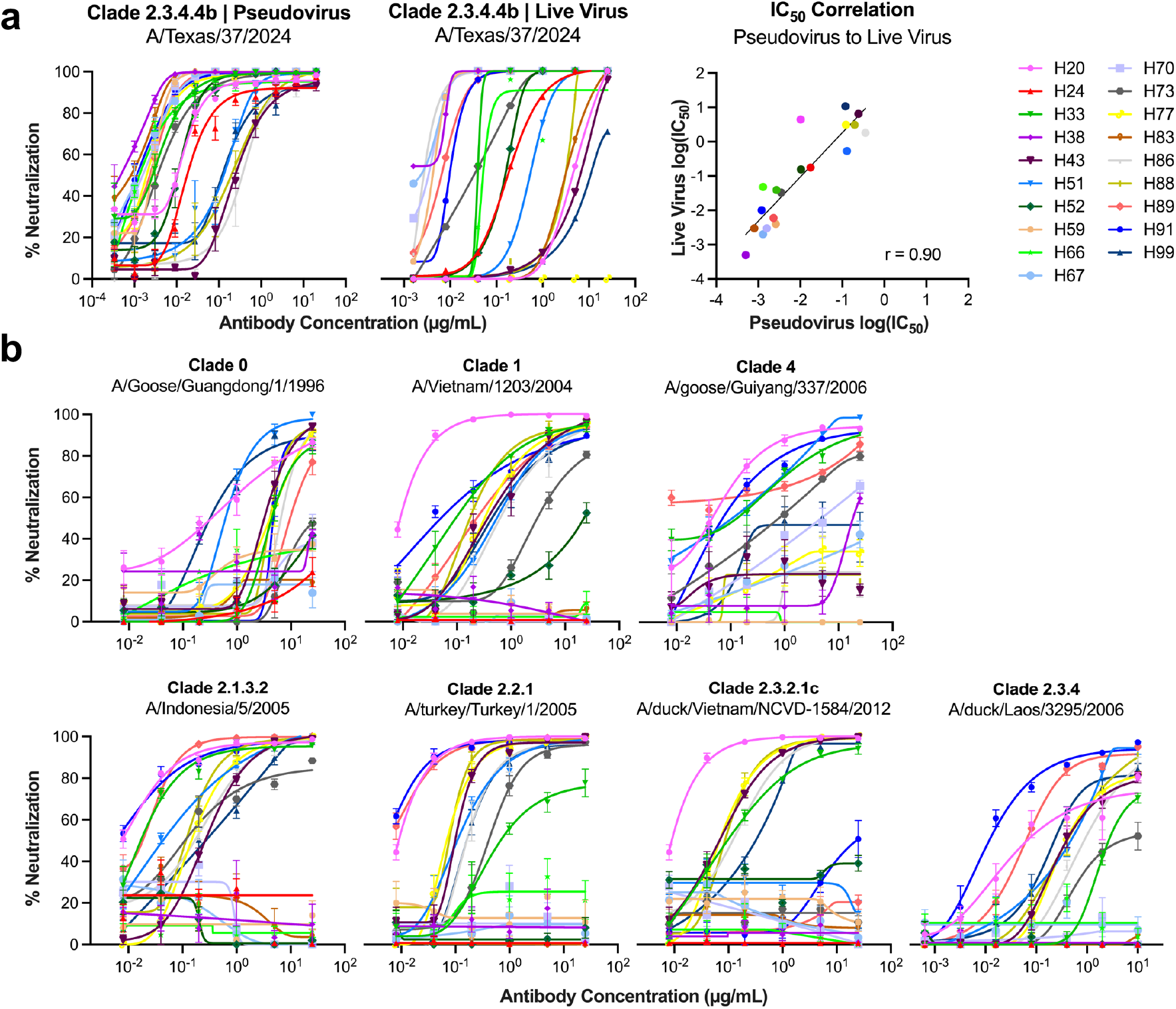
Characterization of neutralizing activities of clade 2.3.4.4b–specific H5 mAbs. a. Neutralization of A/Texas/37/2024 by purified mAbs in pseudovirus (PV) and authentic live-virus (LV) assays, and correlation of IC50 values between PV and LV. b. Neutralization breadth of purified mAbs across a panel of H5 viruses spanning seven clades.

Given the uniformly strong neutralization of the clade 2.3.4.4b strain, we next assessed breadth across the broader global H5 phylogeny. We assembled a PV panel spanning major historical and recent H5 clades indicated in **Figure 1a**, including clades 0, 1, 4, 2.1.3.2, 2.2.1, 2.3.2.1c, 2.3.4, and 2.3.4.4b. Eight mAbs (H24, H38, H52, H59, H66, H67, H70, and H83) were highly specific for the clade 2.3.4.4b lineage and failed to neutralize viruses from earlier or phylogenetically distant H5 clades (**Figure 2b**). The other 11 mAbs displayed varying degrees of cross-clade neutralizing activity, and among these, H91 exhibited exceptional breadth and potency, neutralizing all but one tested strain with IC50 values as low as 1 ng/mL.

### Epitope mapping reveals antibody groups with distinct neutralization patterns

To define the epitope specificities underlying the diversity of H-series antibodies, we performed pairwise competition ELISAs of 17 mAbs in a checkerboard format. Two antibodies, H20 and H43, exhibited weak binding to the HA trimer and were therefore not included in the competition matrix; instead, H20 was characterized structurally by cryo-EM later, whereas H43 was provisionally grouped based on its neutralization profile.

The resulting competition matrix revealed several discrete clusters of antibodies (**Figure 3a**). A tight cluster comprising H38, H67, H59, and H70 showed strong mutual competition and competed with the reference antibody 65C6, which is known to target the vulnerable site 1 (VS1) epitope on the HA head^28^. H73 and H91 formed a largely self-contained competition group, competing strongly with each other and partially with 65C6 but minimally with the H38/H67/H59/H70 cluster, suggesting that they target a head epitope distinct from the VS1 site. In contrast, H83 exhibited minimal competition with other antibodies, revealing a unique epitope among our isolated mAbs.

**Figure 3.**
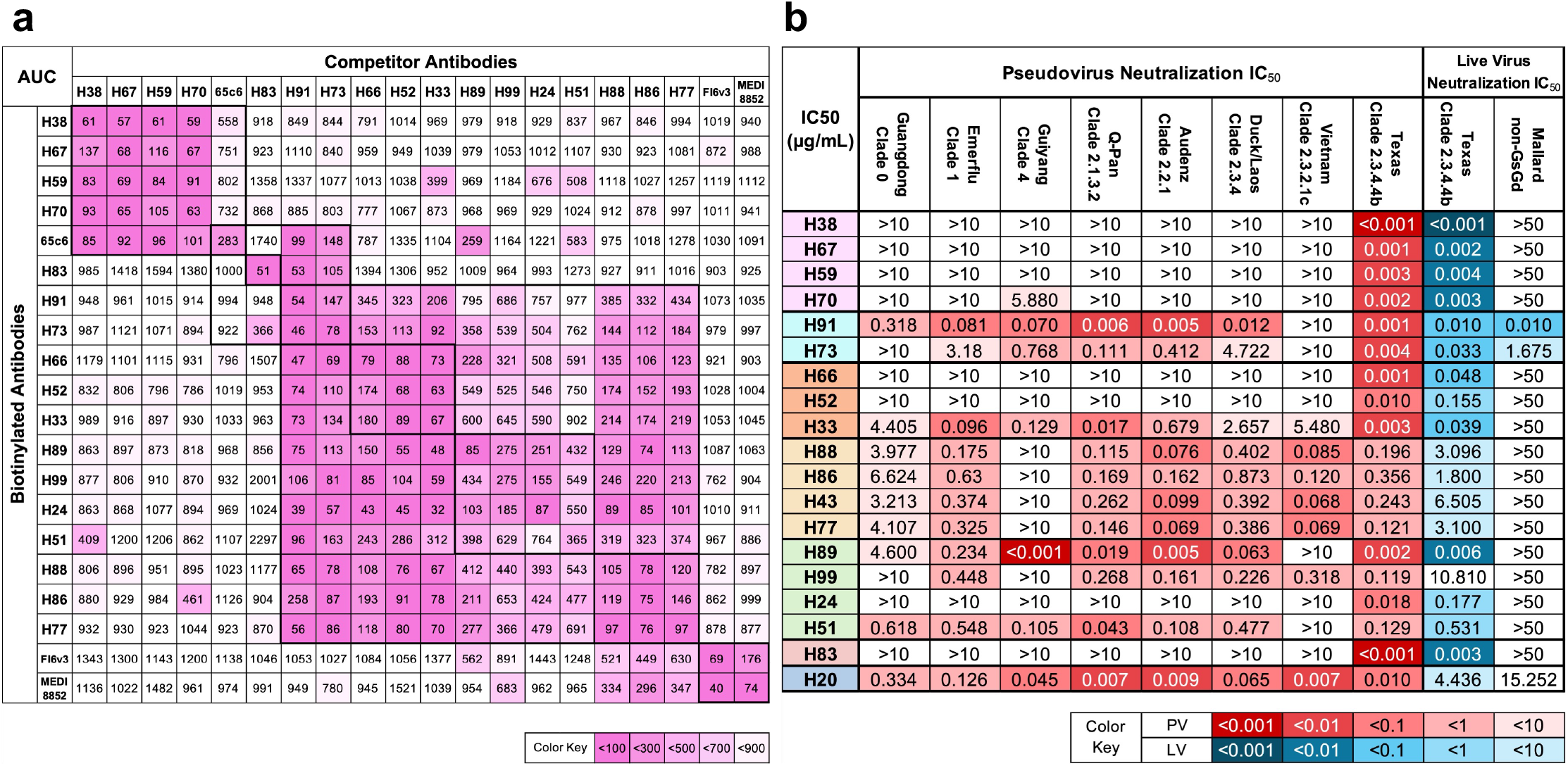
Epitope mapping and neutralization profiles of H5 mAbs. a. Competition ELISA matrix showing the ability of unlabeled HA-binding mAbs to block binding of a biotinylated mAb to the HA trimer. Values denote the area under the competition curve (AUC). b. Neutralization IC50 values for 19 mAbs measured in pseudovirus and live-virus assays, grouped by competition-defined epitope classes.

Beyond these discrete groups, a large cluster of antibodies showed extensive mutual competition but limited competition with the VS1-like group and with H83, H73, and H91, suggesting that they occupy a contiguous yet distinct antigenic surface on HA. The stem-directed antibody controls Fi6v3 and MEDI8852 did not compete with any of the H-series mAbs, confirming that these 17 antibodies are directed to the HA head.

To determine whether these competition-defined groups corresponded to distinct neutralization profiles, we reorganized the IC50 values from the PV neutralization assays (**Figure 2b**) by their epitope clusters (**Figure 3b**). Antibodies in the VS1-like cluster (H38, H67, H59, H70) shared a nearly identical neutralization pattern; they potently neutralized the A/Texas/37/2024 2.3.4.4b virus but showed little or no activity against other clades. In contrast, antibodies belonging to the large central cluster displayed neutralization breadth beyond clade 2.3.4.4b clades, except for H66, H52, and H24. H83 that targets a unique epitope also lacked neutralization breadth, whereas the small cluster of H73 and H91 exhibited exceptional neutralization breadth. Notably, H91 neutralized nearly all strains tested, except for clade 2.3.2.1c; it also neutralized the non-Gs/GD lineage virus A/mallard/Wisconsin/2576/2009 strain, indicating that H91 recognizes a highly conserved epitope shared across phylogenetically diverse H5 viruses (**Figure 3b**).

### Cryo-EM structures reveal epitopes corresponding to antibody groupings

To define the structural basis of antibody recognition, we determined cryo-EM reconstructions of prefusion HA trimer in complex with Fab fragments for mAbs representing each epitope group shown in **Figure 3b** (H70, H83, H91, H33, H77, and H51) plus H20 that could not be tested in the competition assay. Antibody complexes were assembled with a soluble recombinant HA from A/Texas/37/2024, which is the same antigen for which we previously solved a structure in complex with Fab 65C6^35^, and then vitrified for data collection on a Titan Krios. Reconstructions reached overall resolutions of 2.73–3.48 Å (**Figure S2 and Table S2**). Well-resolved Fab density supported atomic model building for H51, H91, H70, H77, H33, and H20. For H83 (3.48 Å resolution) a reconstruction electron density map consisting of a “dimer of trimers” (**Figure S2b**) was obtained with sufficiently resolved cryo-EM density to model the heavy chain in contact with HA. The density for the light chain, which makes minimal contact with HA, was only sufficiently resolved to place the main chain.

The resulting structures revealed a diverse set of HA epitopes that were broadly consistent with the competition-defined clusters (**Figures 4a & 4b**). H70 engaged the VS1 region at the apex of the HA head, with its HCDR3 inserting into a hydrophobic groove; notably, the HCDR3 adopts a binding orientation that mirrors the previously described antibody 20D10^26^, positioning a key tyrosine residue in a nearly superimposable manner (**Figure S3a**). The H83 epitope included the receptor-binding site (RBS) and was consistent with an RBS-directed mode of recognition in which the HCDR3 projects into the pocket and a tryptophan sterically competes with sialylated receptor engagement (**Figure 4c**). In contrast, H51, H91, H77, and H33 all targeted the lateral aspect of the HA head (**Figures 4a & 4b**). Among these, H91 bound higher on the head, spanning the upper lateral patch, whereas H51, H77, and H33 recognized a lower lateral surface; interestingly, all four footprints converge on a shared structural element centered on the “DEFIRVP” motif starting from residue 78 (**Figure 4d**). Finally, H20 bound the HA stem at a site that is distinct from the head-directed antibodies.

**Figure 4.**
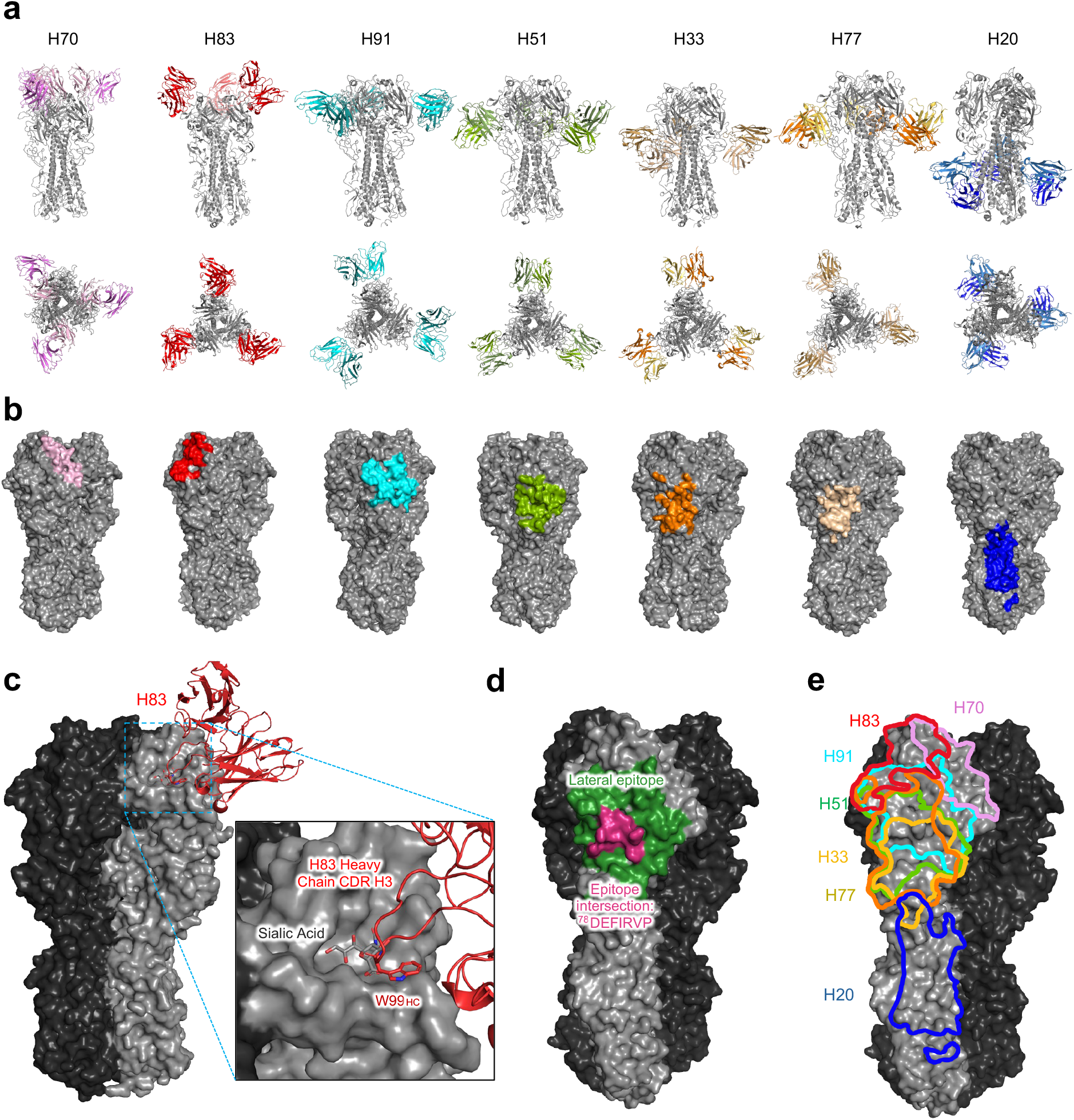
Cryo-EM analysis reveals diverse epitopes recognized by H-series monoclonal antibodies. a. Cryo-EM structures reveal diverse epitopes, modes of recognition, and angles of approach. b. Footprint of each antibody epitope. c. The RBS-directed mAb H83 occupies the receptor-binding site and displaces sialic acid via its CDRH3. d. Intersection of lateral-epitope antibodies (H91, H51, H33, H77). Green indicates the union of residues contacted by any of these mAbs, whereas magenta indicates the shared (intersection) footprint contacted by all four. The common core includes a DEFIRVP motif. e. Footprints of epitopes overlayed on HA

Collectively, projecting these epitope footprints onto the HA trimer showed that the competition groups defined here recognize several major, partially overlapping antigenic regions of the clade 2.3.4.4b HA, spanning the head apex, the receptor-binding site, the upper and lower lateral head surfaces, and the stem (**Figure 4e**). Notably, all mAbs structurally characterized recognized epitopes within a single protomer; none targeted quaternary epitopes spanning across protomers.

### Antibody-defined antigenic differences among recent clade 2.3.4.4b human isolates

Recent human infections in North America were caused primarily by two genetically distinct clade 2.3.4.4b sublineages, the cattle-associated genotype B3.13 and the avian/poultry-associated genotype D1.1. Phylogenetic analysis of available HA sequences revealed ongoing diversification within and between these genotypes (**Figure 5a)**. Relative to the WHO candidate vaccine virus A/American_Wigeon/South_Carolina/AH0195145/2021 (A/American_Wigeon/2021), strain A/Texas/37/2024 acquired L119Q and T199I, and subsequent B3.13 human isolates accumulated additional substitutions including D92G, S324N and V135M. In parallel, D1.1 viruses shared M108L, A214V, and V515I mutations, with additional lineage-specific changes observed in human isolates, including T40A, K329R and N476D in A/Louisiana/12/2024 and A144T in A/British_Columbia/PHL-2032/2024. Notably, the fatal H5N5 infection by A/Washington/2148/2025 carried additional HA substitutions on the D1.1 background, including A160T, which was also observed in the fatal case by A/Mexico/Dur_Indre_2292/2025 (**Figure 5a**). Mapping these substitutions onto the HA trimer showed that most mutations were localized to the HA head (**Figure 5b**), with several directly positioned within or adjacent to the VS1-VS3 epitope clusters (including S124G, E130D, L119Q, V135M, A144T and D240G).

**Figure 5.**
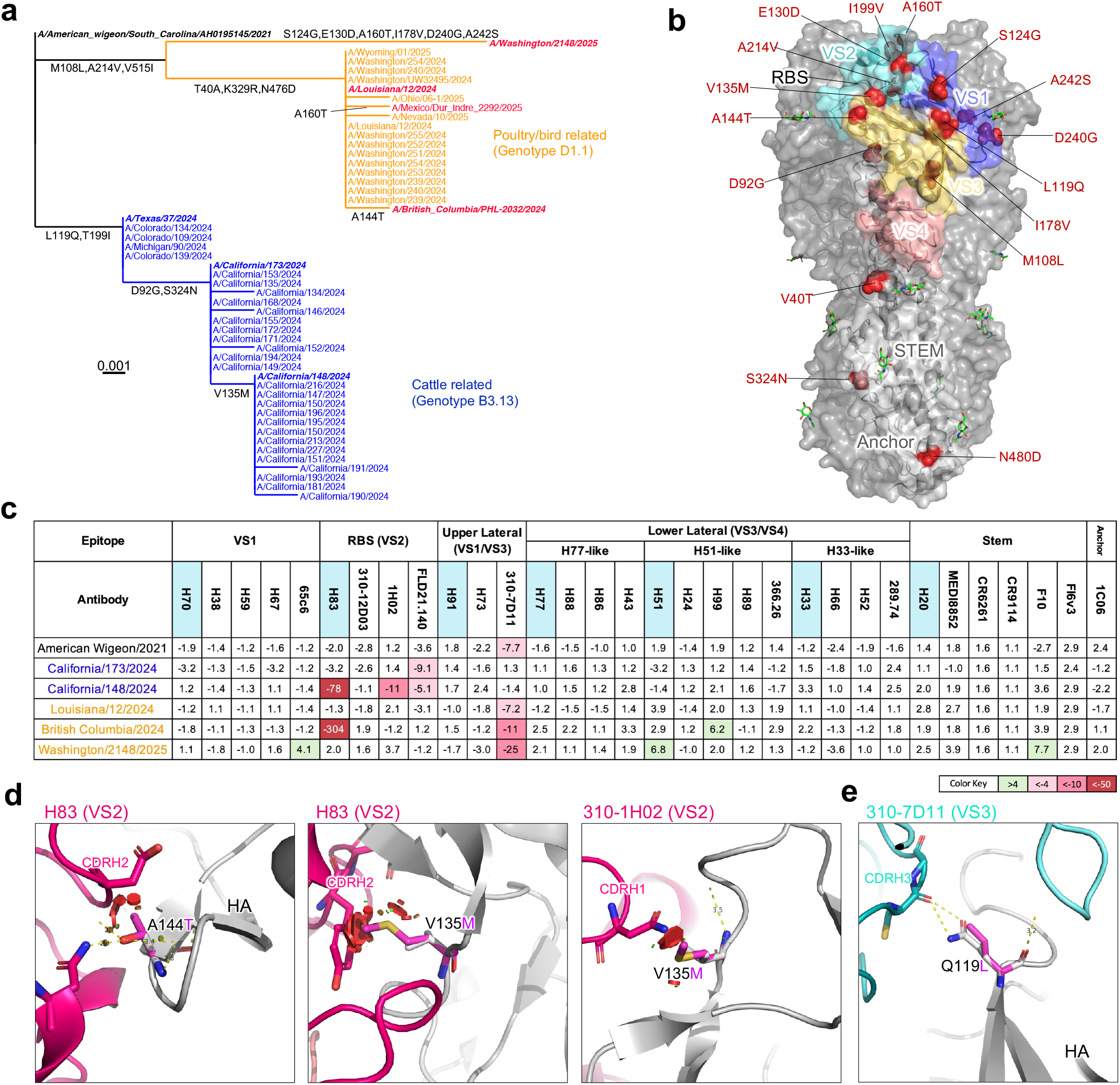
Antigenic tracking of recent clade 2.3.4.4b variants. a. HA phylogenetic tree of recent North American H5Nx human infections. Severe or fatal cases are highlighted in red; variants tested in this study are shown in bold. b. Locations of amino-acid substitutions in recent clade 2.3.4.4b variants mapped onto the HA surface, with major epitope classes indicated. c. Fold change in mAb neutralization IC50 against each variant relative to A/Texas/37/2024. Antibodies without the H prefix are previously published mAbs. d. Structural modeling of A144T and V135M in the context of H83 and 310-1H02 binding; predicted steric clashes are shown as red plates. e. Structural modeling of a substitution of glutamine (Q) with leucine (L) at position 119, observed in ancestral and D1.1 viruses, shows disruptions to the hydrogen-bond interactions with the CDRH3 of 310-7D11.

To determine whether these HA sequence differences were accompanied by changes in antibody susceptibility, we assembled a 32-mAb reference panel comprising our 19 newly isolated H-series mAbs and 13 previously published mAbs with complementary epitope specificities (**Figure S3b**). Together, these antibodies recognize distinct and partially overlapping regions of the HA head (VS1–VS4), stem helix, and anchor, allowing neutralization susceptibility to be compared across multiple antigenic sites. We tested a panel of pseudoviruses representing key recent human clade 2.3.4.4b isolates, including A/American_Wigeon/2021, A/Texas/37/2024, A/California/173/2024, A/California/148/2024, A/Washington/2148/2025, A/Louisiana/12/2024, and A/British_Columbia/PHL-2032/2024 (**Figure 5c**). Using the B3.13 strain A/Texas/37/2024 as the reference, we observed pronounced neutralization escape from the RBS-directed mAb H83 by A/California/148/2024 (B3.13) and A/British_Columbia/PHL-2032/2024 (D1.1), with 78-fold and 304-fold increases in IC50, respectively (**Figure 5c & S4**). In contrast, the upper-lateral HA mAb 310-7D11 exhibited consistent reduction in neutralization potency against A/American_Wigeon/2021 and multiple D1.1 viruses, with ∼7-25-fold increases in IC50 values (**Figure 5c**), indicating incremental antigenic drift within this otherwise conserved epitope.

To investigate the molecular basis of antibody escape, we modeled antibody-HA complexes using available structural templates. The V135M and A144T, located at the periphery of the RBS, were predicted to introduce steric clashes that could interfere with H83 and 310-1H02 binding (**Figure 5d**). For the upper-lateral antibody 310-7D11, modeling suggested that the glutamine (Q) at position 119 in B3.13 viruses forms a hydrogen-bonding interaction with the antibody CDRH3, whereas the leucine (L) present in ancestral and D1.1 viruses lacks this interaction (**Figure 5e**). These structural predictions were consistent with the observed differences in neutralization susceptibility and prompted further investigation of the individual substitutions.

To assess whether these substitutions individually affect antibody recognition, we introduced V135M, A144T, and Q119L into the A/Texas/37/2024 HA background and expressed and purified the corresponding recombinant trimers. By ELISA, A144T abolished and V135M reduced H83 binding, whereas Q119L reduced binding of 310-7D11. Binding of H38, which served as a control, was unaffected by these substitutions (**Figure S4**). In addition to substitutions potentially affecting direct antibody contacts, we investigated A160T, which was identified in two recent fatal human isolates and introduces a potential N-linked glycosylation site at Asn158 near the RBS (**Figures S4a & b**). Because glycan acquisition can alter the accessibility of HA antigenic surfaces, we examined the electrophoretic mobility of recombinant HA carrying A160T. The mutant HA migrated more slowly than wild-type HA by SDS-PAGE (**Figure S4c**), consistent with the possibility of an addition of an N-linked glycan.

## Discussion

The continued diversification of clade 2.3.4.4b H5N1 viruses underscores the need to characterize the antigenic consequences of emerging HA mutations. Here, we isolated 19 human mAbs with diverse neutralization profiles and epitope specificities, including antibodies with broad activity against phylogenetically distinct H5 viruses. By combining these antibodies with previously characterized mAbs, we established a 32-mAb reference panel that serves not only as a collection of potent neutralizing antibodies but also as a tool for characterizing antigenic changes at the molecular level. This structurally informed, epitope-resolved panel enables systematic assessment of how naturally occurring HA substitutions affect antibody recognition across circulating H5N1 viruses.

A notable finding of our study is the identification of broadly neutralizing antibodies targeting the lateral surface of the HA head. Structural analyses revealed several partially overlapping epitopes in this region, including a conserved structural element containing the DEFIRVP motif. Among these antibodies, H91 exhibited exceptional breadth and potency, neutralizing nearly all H5 strains tested. These findings identify the lateral HA head as an important site of vulnerability, where antibodies can achieve potent neutralization while recognizing surfaces conserved across divergent H5 lineages. Monitoring mutations within these regions may therefore provide valuable insights into emerging patterns of antibody escape.

Application of our antibody panel to recent North American human isolates revealed that antibody-defined antigenic differences are already detectable among clade 2.3.4.4b viruses, with distinct patterns of escape observed across B3.13 and D1.1 isolates. We observed pronounced reductions in susceptibility to RBS-directed antibodies associated with substitutions near the RBS, including V135M and A144T. Structural modeling suggested that these substitutions could interfere with antibody binding, while mutant HA experiments demonstrated reduced or abolished recognition by H83. We also identified A160T in two recent isolates, which introduces a potential N-linked glycosylation site near the RBS. The observed electrophoretic mobility shift of A160T HA was consistent with glycan addition, providing a plausible additional mechanism of escape through glycan shielding. In contrast, the upper-lateral neutralizing antibody 310-7D11 exhibited reduced neutralization that was associated with a leucine (L) at position 119. This residue is present in the D1.1 and ancestral viruses, but not in B3.13 viruses. These findings reveal distinct, genotype-specific patterns of antibody escape across the HA head, ranging from pronounced reductions in susceptibility to RBS-directed antibodies to more moderate changes affecting conserved lateral-head epitopes.

Together, our findings establish a structurally informed antibody reference panel for characterizing the ongoing antigenic evolution of H5 viruses. By integrating neutralization profiles with competition-defined epitope groups and cryo-EM structures, this panel provides a framework for interpreting how naturally occurring HA substitutions affect antibody recognition across distinct antigenic regions. Coupled with genomic surveillance, such analyses could complement existing approaches, including antigenic cartography and deep mutational scanning, to inform the evaluation of circulating H5 viruses and the development of vaccines and antibody-based countermeasures. Importantly, this approach enables characterization of antigenic changes in circulating viruses rather than prediction of future evolutionary trajectories, which are also influenced by viral fitness, receptor binding, and transmissibility. In addition, broadly neutralizing antibodies such as H91 may warrant further investigation for prophylactic or therapeutic applications.

### Limitations of the study

Several features of the experimental system define the scope of the conclusions drawn here. The panel derives from a defined immunization regimen in VelocImmune mice, and the epitopes it samples reflect that regimen together with the immunodominance hierarchy of this model, which may differ from that of human H5 exposure or vaccination. Epitope assignments rest on competition profiles for 17 antibodies and cryo-EM structures for 7; they therefore resolve the antigenic regions represented within the panel, and other surfaces of HA may carry antigenic variation that these antibodies do not report. Antigenic comparisons were made predominantly in a pseudovirus system, which measures entry inhibition, with authentic-virus measurements for two strains. The viruses analyzed were drawn from recent North American human infections and define the temporal, geographic and host range over which the antigenic relationships described here apply. The contribution of individual substitutions was resolved at the level of HA binding. For A160T, the mobility of the mutant protein is consistent with an added N-linked glycan; the extent of glycan occupancy and its consequences for antibody recognition remain to be defined. The measurements reported here describe recognition by individual mAbs. Relating them to population-level immunity or to vaccine performance requires additional studied using polyclonal sera. The antibodies have been characterized in vitro, and their activity in vivo has not been examined

## STAR METHODS

Detailed methods are provided in the online version of this paper and include the following:

### RESOURCE AVAILABILITY

#### Lead contact

Further information and requests for resources and reagents should be directed to and will be fulfilled by the Lead Contact Author David D. Ho.

#### Materials availability

All reagents generated in this study are available from the Lead Contact with a completed Materials Transfer Agreement.

#### Data and code availability

Data reported in this paper will be shared by the lead contact upon request. Cryo-EM maps for HA complexes with H70 (EMD-74842), H83 (EMD-74879), H91 (EMD-74755), H51 (EMD-74844), H33 (EMD-74873), H77 (EMD-74865), and H20 (EMD-74855) have been deposited to the EMDB., Fitted coordinates for H70 (9ZV0), H83 (9ZVL), H91 (9ZTJ), H51 (9ZV1), H33 (9ZVH), H77 (9ZV9), and H20 (9ZV4) have been deposited to PDB. The sequences encoding the variable regions of all 19 H-series monoclonal antibodies have been deposited in GenBank under accession numbers QB248878-QB248915. Any additional information required to reanalyze the data reported in this paper is available from the lead contact upon request.

### EXPERIMENTAL MODEL AND SUBJECT DETAILS

#### Cell lines

HEK293T (CRL-3216) and MDCK (CCL-34) cells were purchased from ATCC and cultured at 37°C with 5% CO2 in Dulbecco’s Modified Eagle Medium (DMEM) + 10% fetal bovine serum (FBS)

+ 1% penicillin-streptomycin. Expi293 cells (A14527) were purchased from Thermo Fisher Scientific and maintained in Expi293 expression medium per the manufacturer’s instructions. The morphology of each cell line was confirmed visually before use. All cell lines tested mycoplasma negative. HEK293T, MDCK, and Expi293 cells are of female origin.

### METHOD DETAILS

#### Plasmid generation

Codon-optimized genes encoding the hemagglutinin (HA) from A/goose/Guangdong/1/1996 (clade 0), A/Vietnam/1203/2004 (clade 1), A/goose/Guiyang/337/2006 (clade 4), A/chicken/Vietnam/NCVD-03/2008 (clade 7.1), A/Indonesia/5/2005 (clade 2.1.3.2), A/turkey/Turkey/1/2005 (clade 2.2.1), A/duck/Vietnam/NCVD-1584/2012 (clade 2.3.2.1c), A/duck/Laos/3295/2006 (clade 2.3.4), and A/Texas/37/2024 (clade 2.3.4.4b) were synthesized and cloned into the pTwist-CMV mammalian expression vector (Twist Biosciences) for pseudovirus production. Individual HA substitutions (V135M, A144T, Q119L, and A160T) were introduced into the A/Texas/37/2024 HA expression construct following the Site-Directed Mutagenesis Kit per manufacturer’s instruction (Agilent) for subsequent characterization. For soluble HA trimers, the ectodomains of the HA were subcloned into the p3BNC mammalian expression vector, followed by a FoldOn trimerization domain and either a 6x His tag, FLAG, myc tag, Strep-tag, or Avi tag. For the generation of B-cell sorting probes, the Y98F mutation was introduced to reduce binding to sialic acid receptors.

#### Protein expression and purification

Plasmids were transfected into Expi293 cells using 1 mg/mL PEI-MAX, and supernatants were collected after five days. Proteins were purified with Strep-Tactin^®^XT 4Flow^®^ (IBA Lifesciences) following the manufacturer’s instructions. Molecular weight and purity were confirmed by SDS-PAGE protein electrophoresis and size-exclusion chromatography prior to use.

#### Mouse vaccination

VelocImmune^®^ female mice were immunized with 10 µg each of plasmids encoding the H5N1 hemagglutinin (HA) from clade 0, clade 7.1, clade 2.2.1, and clade 2.3.4.4b, delivered into separate limbs by DNA electroporation at weeks 0 and 3. Protein boosts (10 µg of purified soluble HA trimer per dose) were administered intramuscularly at weeks 5, 8, and 20. Peripheral blood was collected two weeks after each immunization to monitor serum ELISA binding and neutralization titers. Splenocytes and peripheral blood mononuclear cells (PBMCs) were isolated at week 30 for B-cell sorting and downstream antibody characterization. All animal procedures were reviewed and approved by the Institutional Animal Care and Use Committee of Columbia University.

#### Sorting for hemagglutinin-specific B cells and single-cell B cell receptor sequencing

Splenocytes and PBMCs were stained with the LIVE/DEAD™ Fixable Yellow Dead Cell Stain Kit (Invitrogen) at room temperature for 10 min, washed with RPMI-1640 complete medium supplemented with 2% FBS, and incubated with 10 µg/ml of clade 0 HA trimer containing a Myc tag and clade 2.3.4.4b HA trimer containing a FLAG tag at room temperature for 45 min. Cells were washed again and stained at 4°C for 45 min with a cocktail of flow-cytometry antibodies, including CD3 PerCP-Cy5.5 (BioLegend), CD19 PE (BioLegend), B220 APC-H7 (BD Biosciences), anti-FLAG BV421 (BioLegend), and anti-Myc APC (BioLegend). Stained cells were washed, resuspended in RPMI-1640 + 2% FBS, and sorted for HA-specific memory B cells (CD3⁻ CD19⁺ B220⁺ HA trimer⁺ live single lymphocytes).

Sorted cells were loaded onto a Chromium X controller using the 5′ Single Cell Immune Profiling Assay (10x Genomics) at the Columbia University Genome Center. Library preparation and quality-control steps were performed according to the manufacturer’s instructions, and sequencing was carried out on an Illumina NextSeq 500 platform.

#### Single cell gene expression analysis and identification of HA specific antibody transcripts

The assembly of full-length antibody and other gene transcripts was performed utilizing the Cell Ranger V(D)J analysis software (version 3.1.0, 10x Genomics), employing default settings with the GRCh38 genome and V(D)J germline sequence version 2.0.0 as the reference. To determine the cell type of B cells, we utilized Seurat package version 4.1.0. Cells with fewer than 500 unique molecular identifiers (UMIs), more than 4500 UMIs, or expression mitochondria gene fraction of more than 5% were discarded in the following analysis. The UMI counts were normalized to 10,000 counts per cell and converted to log scale (Seurat function NormalizeData). nUMI were regressed out using the ScaleData function. Highly variable genes (n=3000) were computed and used as input for principal components analysis. Significant principal components were used for downstream graph-based, semi-unsupervised clustering into distinct populations (FindClusters function), and Uniform Manifold Approximation and Projection (UMAP) dimensionality reduction was then used to project these populations in two dimensions. For clustering, the resolution parameter, which indirectly controls the number of clusters, was approximated based on the number of cells according to Seurat guidelines; a vector of resolution parameters was passed to the FindClusters function, and the optimal resolution that established discernible clusters with distinct marker gene expression was selected. To identify marker genes, the clusters were compared pairwise for differential gene expression using the Wilcoxon rank sum test for single-cell gene expression (FindAllMarkers function, min.pct = 0.4, min.diff.pct = 0.3, and false discovery rate < 0.05). To assign identities to these subpopulations, we cross-referenced their marker genes with known B cell subtype markers from the literature, Fcer2a for Naïve B cell, Cd80 for memory B cell, Itgax for atypical memory B, Sdc1 and Xbp1 for plasma blast.

#### Antibody transcript annotation

Transcripts specific to the antigen were analyzed and annotated with SONAR version 2.0, following previously established procedures^32^. Assignment of V(D)J gene segments to each transcript was conducted via BLASTn, employing specialized parameters against a germline gene repository sourced from the International ImMunoGeneTics (IMGT) information system database. The identification of the complementarity-determining region 3 (CDR3) utilized BLAST alignments of the V and J segments, focusing on the conserved second cysteine within the V segment and the WGXG (for heavy chains) or FGXG (for light chains) motifs in the J segment, with “X” indicating any amino acid. Isotype determination for heavy chain transcripts was achieved by analyzing Constant domain 1 (CH1) sequences against a human CH1 gene database from IMGT, using BLASTn with standard parameters. The CH1 allele presenting the lowest E-value was selected for precise isotype classification, adhering to a BLAST E-value cutoff of 10e-6. Transcripts with incomplete V(D)J segments, frameshifts, or extraneous sequences beyond the V(D)J region were discarded. The filtered transcripts were then aligned to their corresponding germline V gene using CLUSTALO, and levels of somatic hypermutation were quantified through the Sievers method. In instances where cells possessed multiple high-quality heavy or light chains, potentially indicative of doublets, combinations of all H and L chains were generated.

#### Antibody expression and purification

For each antibody, variable region genes were codon-optimized for human cell expression and synthesized by Twist Bioscience. The VH and VL genes were cloned separately into gWiz mammalian expression vectors encoding the respective human IgG heavy- and light-chain constant regions. Monoclonal antibodies were expressed in Expi293 cells by co-transfection of the heavy- and light-chain plasmids using PEI-MAX (1 mg/mL) and cultured at 37°C shaking at 125 rpm in 8% CO2. At day 3 post-transfection, culture supernatant was collected for screening antibody binding by ELISA, and at day 4, additional supernatant was collected for pseudovirus neutralization assays. The remaining culture supernatants were harvested on day 5 for purification using rProtein A Sepharose (Cytiva) affinity chromatography per the manufacturer’s instructions.

#### ELISA

High-binding 96-well plates (Corning Costar) were coated with 50 ng per well of HA trimer in PBS and incubated overnight at 4°C. After washing with PBST (0.05% Tween-20 in PBS), plates were blocked with 200 µL of blocking buffer (1% BSA in PBS) at 37°C for 2 h. Plates were washed again before adding antibody-containing supernatants or purified monoclonal antibodies, which were serially diluted in blocking buffer and incubated for 1 h at 37°C. After washing, 100 µL of horseradish peroxidase (HRP)-conjugated goat anti-human IgG (H+L) secondary antibody (Jackson ImmunoResearch; 1:5,000 dilution) was added and incubated for 1 h at 37°C. Following a final wash, Ultra TMB-ELISA substrate solution (ThermoFisher) was added, and the reaction was stopped with 1M sulfuric acid. Absorbance was measured at 450 nm using a microplate reader. All ELISAs were performed in technical duplicates or triplicates, with selected antibody– antigen combinations independently repeated to confirm reproducibility.

#### Competition ELISA

Purified monoclonal antibodies were biotinylated using the One-Step Antibody Biotinylation Kit (Miltenyi Biotec) according to the manufacturer’s instructions and subsequently purified with a 50 kDa Amicon^®^ Ultra centrifugal filter (Sigma). Serially diluted competitor antibodies (50 µL per well) were added to HA trimer-coated ELISA plates, followed by 50 µL of biotinylated antibodies at a concentration that yielded an OD450 value of approximately 1.0 in the absence of competitor antibodies. Plates were incubated at 37°C for 1 h, washed with PBST (0.05% Tween-20 in PBS), and then incubated for another 1 h at 37°C with 100 µL of Pierce™ High Sensitivity NeutrAvidin™-HRP (ThermoFisher) diluted 1:10,000. Plates were washed again before adding Ultra TMB-ELISA substrate solution (ThermoFisher), and the reaction was stopped with 1M sulfuric acid. Absorbance was measured at 450 nm using a microplate reader. For all competition ELISA experiments, the relative binding of biotinylated antibodies to the HA trimer in the presence of competitors was normalized to the signal obtained in competitor-free wells. Relative binding curves and area under the curve (AUC) values were determined by fitting non-linear five-parameter logistic (5PL) dose-response models in GraphPad Prism v11.

#### Pseudovirus production and infectivity

HEK293T cells were co-transfected with a HA-expressing and NA-expressing construct using 1 mg/mL PEI-MAX and then infected with VSV-G pseudotyped ΔG-luciferase (G*ΔG-luciferase, Kerafast) 1-day post-transfection. Two hours after infection, cells were washed three times with PBS, changed to fresh medium, and then cultured for one more day before the cell supernatants were harvested. Anti-VSVG (anti-I1) antibody was added to deplete non-pseudotyped viruses. Pseudoviruses were then harvested, centrifuged, and then aliquoted and stored at -80°C.

#### Pseudovirus neutralization assays with sera or mAbs

Each H5N1 pseudovirus was titrated to standardize viral infectious dose before use in neutralization assays. Seven serial dilutions of heat-inactivated mouse sera, or monoclonal antibodies (mAbs) were added in 96-well plates, starting at 1:50 dilution for sera and 10 µg/mL for antibodies. Next, pseudoviruses were added and incubated at 37°C for 1 hour. In each plate, wells containing only pseudoviruses were included as controls. 3×10^4^ MDCK cells were then added per well and incubated at 37°C for 20 hours. Promega Luciferase Assay System (E4550) was used for lysis and luciferase activity measurements on a Tecan Infinite^®^ 200 PRO using i-control™ software v.3.9.1.0, in accordance with the manufacturer’s instructions. The serum dilution or mAb concentration that inhibits 50% of virus entry (ID50 or IC50) was calculated using nonlinear five-parameter dose-response curve fitting using GraphPad Prism v.11. Neutralization assays were performed in technical triplicates, with selected antibodies and viral strains independently retested to confirm reproducibility.

#### Infectious H5N1 neutralization assays

Authentic live virus neutralization assay was performed as previously described^18^. Highly pathogenic influenza virus A/bovine/Ohio/B24OSU-439/2024 (H5N1) and low-pathogenicity influenza virus A/mallard/Wisconsin/2576/2009 (H5N1) were obtained from BEI Resources. The HA amino acid sequence of A/bovine/Ohio/B24OSU-439/2024 is identical to that of A/Texas/37/2024, which was used in the pseudovirus neutralization assays. Viruses were propagated in MDCK cells at 37°C with 5% CO₂ for 4 days before titration. Monoclonal antibodies were serially diluted fivefold in DMEM supplemented with 10% FBS, starting at 50 µg/mL. Each antibody dilution was incubated with 40 TCID50 of H5N1 virus at 37°C for 30 min, followed by addition of 100 µL of the antibody-virus mixture to confluent MDCK cell monolayers seeded the previous day. Plates were incubated at 37°C with 5% CO2 for 96 h, after which cytopathic effects (CPE) were visually assessed. CPE in each well was scored relative to virus-only control wells to determine percent neutralization. The antibody concentration required to inhibit 50% of virus infection (IC50) was calculated by fitting non-linear five-parameter logistic (5PL) dose-response curves using GraphPad Prism v11.

#### HA Construct Design and Protein Production for Cryo-EM

The sequence of H5N1 HA (A/Texas/37/2024) was expressed as previously described^35^ to replace the Furin cleavage site (REKRRKR) with (RRRRRR) for enhanced Furin cleavage. The transmembrane and cytosolic regions were removed and replaced with a 2X GSS linker, followed by a T4 Fold domain (GYIPEAPRDGQAYVRKDGEWVLLSTFL), glycine residue, 10x HIS tag, GGSG linker, and then AVI tag. Codon optimized DNA was synthesized by Genscript and subcloned into the gWiz vector backbone for transient expression in mammalian cells. Soluble, fully cleaved H5 HA trimers were produced by transient co-transfection with Furin enzyme in Expi293F cells (Life Technologies) using Expi293 Transfection reagent. After 5 days at 37 °C, culture supernatants were harvested by centrifugation. The recombinant HA trimer was captured by Ni-NTA (Sigma-Aldrich). The resin was washed with 1X PBS, pH 7.5, 25 mM imidazole, and then protein was eluted with 1X PBS, pH 7.5, 550 mM imidazole. The eluant was purified by size exclusion chromatography on a Superdex 200 Increase 10/300 GL column (Cytiva) into 1X PBS, pH 7.5.

#### Cryo-EM Grid Preparation

Samples for cryo-EM grid preparation were produced by first mixing 11 µl of purified H5N1 HA (A/Texas/37/2024) at 1.1 mg/ml with 8 µl of Fab ranging from 1-3 mg/ml. Complex was adjusted to have a final concentration of 0.005% (w/v) n-Dodecyl β-D-maltoside (DDM) to prevent preferred orientation and aggregation during vitrification and incubated on ice for 20 minutes. Cryo-EM grids were prepared by applying 3 μL of sample to a freshly glow discharged carbon-coated copper grid (CF 1.2/1.3 300 mesh). The sample was vitrified in liquid ethane using a Vitrobot Mark IV with a wait time of 30 s, a blot time of 3 s, and a blot force of -5.

#### Image Processing

Cryo-EM data were collected on a Titan Krios operating at 300 keV, equipped with a K3 detector (Gatan) operating in counting mode. Data were acquired using EPU (ThermoFisher). The dose was fractionated over 50 raw frames. For all structures, the movie frames were aligned and dose-weighted using cryoSPARC 4.7.1; the CTF estimation, particle picking, 2D classifications, ab initio model generation, heterogeneous refinements, homogeneous 3D refinements and non-uniform refinement calculations were carried out using cryoSPARC 4.7.1.

#### Atomic Model Building and Refinement

For structural determination, a starting model of the antibody Fab was obtained using SABPRED^36^. For HA, the previously published cryoEM structure of H5 influenza hemagglutinin from clade A/Texas/37/2024 (PDB 9EKF) was used. The Fab and HA starting models were docked into the cryo-EM density map using UCSF Chimera to build an initial model of the complex. The model was then manually rebuilt to the best fit into the density using Coot^37^ and refined using Phenix36. Interface calculations were performed using PISA. Structures were analyzed and figures were generated using PyMOL (http://www.pymol.org) and UCSF Chimera. Final model statistics are summarized in **Figure S2**.

### QUANTIFICATION AND STATISTICAL ANAYLYSIS

Neutralization ID50 and IC50 values were determined by fitting a five-parameter dose-response curve in GraphPad Prism v11.

#### Author Contributions

Conceptualization, Y.G., D.D.H.; supervision, management, and funding acquisition, Y.H., P.D.K, Y.G., and D.D.H.; investigation, data curation, and analysis, all authors; writing, review, and editing, H.H., N.C.M., M.W., S.C., Y.H., P.D.K., Y.G., and D.D.H.

## Acknowledgements

This study was supported by funding from the Gates Foundation (INV019355) and donations from Andrew and Peggy Cherng, Roger and David Wu, and Mark Kingdon & Anla Cheng. The VelocImmune mice were kindly provided by Regeneron Pharmaceuticals.

---

**Table S1.** Neutralization potency and binding affinity of published mAbs against H5N1 clade 0 and clade 2.3.4.4b viruses. NB indicates not binding in SPR.

| **Epitope** | **mAbs** | **SPR KD (M)** |  | **ELISA EC<sub>50</sub> (µg/mL)** |  | **Neutralization IC<sub>50</sub> (µg/mL)** |  |
|---|---|---|---|---|---|---|---|
|  |  | **Clade 0** | **Clade 2.3.4.4b** | **Clade 0** | **Clade 2.3.4.4b** | **Clade 0** | **Clade 2.3.4.4b** |
| VS1 | 3C11 | 5.0E-11 | 4.6E-09 | 0.004 | 0.008 | 0.470 | >10 |
|  | 65C6 | 1.5E-10 | 4.3E-09 | 0.005 | 0.012 | 0.027 | 0.075 |
|  | AVFluIgG01 | 1.1E-10 | NB | 0.007 | >10 | 0.050 | >10 |
|  | FLD194 | 2.6E-10 | NB | 0.004 | >10 | 0.012 | >10 |
| VS2 | FLD21.140 | 1.2E-08 | 1.9E-08 | >10 | 9.691 | 0.725 | 0.501 |
| VS4 | H5M9 | NB | NB | >10 | >10 | >10 | >10 |
| VS5 | FluA-20 | 1.2E-08 | NB | 1.424 | >10 | >10 | >10 |
| Stem | CR6261 | 6.9E-10 | 8.2E-10 | 0.260 | 0.013 | 0.011 | 0.013 |
|  | CR9114 | 2.0E-11 | 2.9E-10 | 0.009 | 0.009 | 0.032 | 0.015 |
|  | F10 | 1.8E-10 | 1.8E-10 | 0.043 | 0.030 | 0.030 | 0.033 |
|  | FI6v3 | 3.1E-13 | 3.0E-13 | 0.009 | 0.008 | 0.028 | 0.012 |
| Anchor | 4F04 | 8.4E-07 | 4.2E-07 | >10 | >10 | >10 | >10 |
|  | 1C06 | 2.4E-07 | 3.4E-06 | >10 | >10 | 9.044 | >10 |
|  | 2F04 | 2.3E-07 | 4.2E-08 | >10 | >10 | >10 | >10 |

**Clade 0**: A/Goose/Guangdong/1/1996
**Clade 2.3.4.4b**: A/American\_wigeon/South\_Carolina/AH0195145/2021

**Table S2.** CryoEM information

| Antibody | H70 | H83 | H91 | H51 | H33 | H77 | H20 |
|---|---|---|---|---|---|---|---|
| EMDB ID | EMD-74842 | EMD-74879 | EMD-74755 | EMD-74844 | EMD-74873 | EMD-74865 | EMD-74855 |
| PDB ID | PDB 9ZV0 | PDB 9ZVL | PDB 9ZTJ | PDB 9ZV1 | PDB 9ZVH | PDB 9ZV9 | PDB 9ZV4 |
| Data collection and processing |  |  |  |  |  |  |  |
| Magnification | 105000 | 105000 | 105000 | 105000 | 105000 | 105000 | 105000 |
| Voltage (kV) | 300 | 300 | 300 | 300 | 300 | 300 | 300 |
| Electron exposure (e–/Å2) | 58 | 58 | 58 | 58 | 58 | 58 | 58 |
| Defocus range (μm) | .8-2 | .8-2 | .8-2 | .8-2 | .8-2 | .8-2 | .8-2 |
| Pixel size (Å) | 0.825 | 0.825 | 0.825 | 0.825 | 0.825 | 0.825 | 0.825 |
| Symmetry imposed | C3 | C3 | C3 | C3 | C3 | C3 | C3 |
| Initial particle images (no.) | 307715 | 80159 | 175085 | 243417 | 209713 | 287561 | 526208 |
| Final particle images (no.) | 228102 | 30030 | 105159 | 183858 | 201510 | 241526 | 259666 |
| Map resolution (Å) | 3.24 | 3.48 | 3.08 | 2.73 | 2.93 | 2.79 | 2.80 |
| FSC threshold | 0.143 | 0.143 | 0.143 | 0.143 | 0.143 | 0.143 | 0.143 |
| Refinement |  |  |  |  |  |  |  |
| Initial model used (PDB code) | 9EKF | 9EKF | 9EKF | 9EKF | 9EKF | 9EKF | 9EKF |
| Model resolution (Å) | 3.24 | 3.48 | 3.08 | 2.7 | 2.93 | 2.79 | 2.80 |
| FSC threshold | 0.143 | 0.143 | 0.143 | 0.143 | 0.143 | 0.143 | 0.143 |
| Model composition |  |  |  |  |  |  |  |
| Non-hydrogen atoms | 17561 | 17372 | 17531 | 17300 | 17141 | 17756 | 17456 |
| Protein residues | 2160 | 2169 | 2145 | 2133 | 2115 | 2166 | 2154 |
| Glycan residues | 33 | 12 | 33 | 33 | 30 | 42 | 33 |
| B factors (Å2) |  |  |  |  |  |  |  |
| Protein | 59.55 | 60.5 | 60.76 | 60.07 | 60.51 | 60.12 | 60.76 |
| Glycans | 30.63 | 79.81 | 49.1 | 30.63 | 30.69 | 30.51 | 45.46 |
| R.m.s. deviations |  |  |  |  |  |  |  |
| Bond lengths (Å) | 0.013 | 0.004 | 0.01 | 0.006 | 0.004 | 0.006 | 0.006 |
| Bond angles (°) | 1.19 | 0.888 | 1.196 | 1.208 | 0.848 | 1.229 | 0.974 |
| Validation |  |  |  |  |  |  |  |
| MolProbity score | 1.99 | 1.94 | 1.9 | 1.93 | 1.9 | 1.78 | 1.94 |
| Clashscore | 4.2 | 3.61 | 6.17 | 4.07 | 3.49 | 4.91 | 5.23 |
| Poor rotamers (%) | 2.96 | 2.55 | 2.36 | 2.33 | 2.77 | 1.78 | 3.32 |
| Ramachandran plot |  |  |  |  |  |  |  |
| Favored (%) | 93.43 | 92.21 | 95.99 | 92.84 | 93.62 | 95.19 | 92.25 |
| Allowed (%) | 6.15 | 7.27 | 3.87 | 6.4 | 6.09 | 4.67 | 7.61 |
| Disallowed (%) | 0.42 | 0.51 | 0.14 | 0.76 | 0.29 | 0.14 | 0.14 |

**Figure S1.**
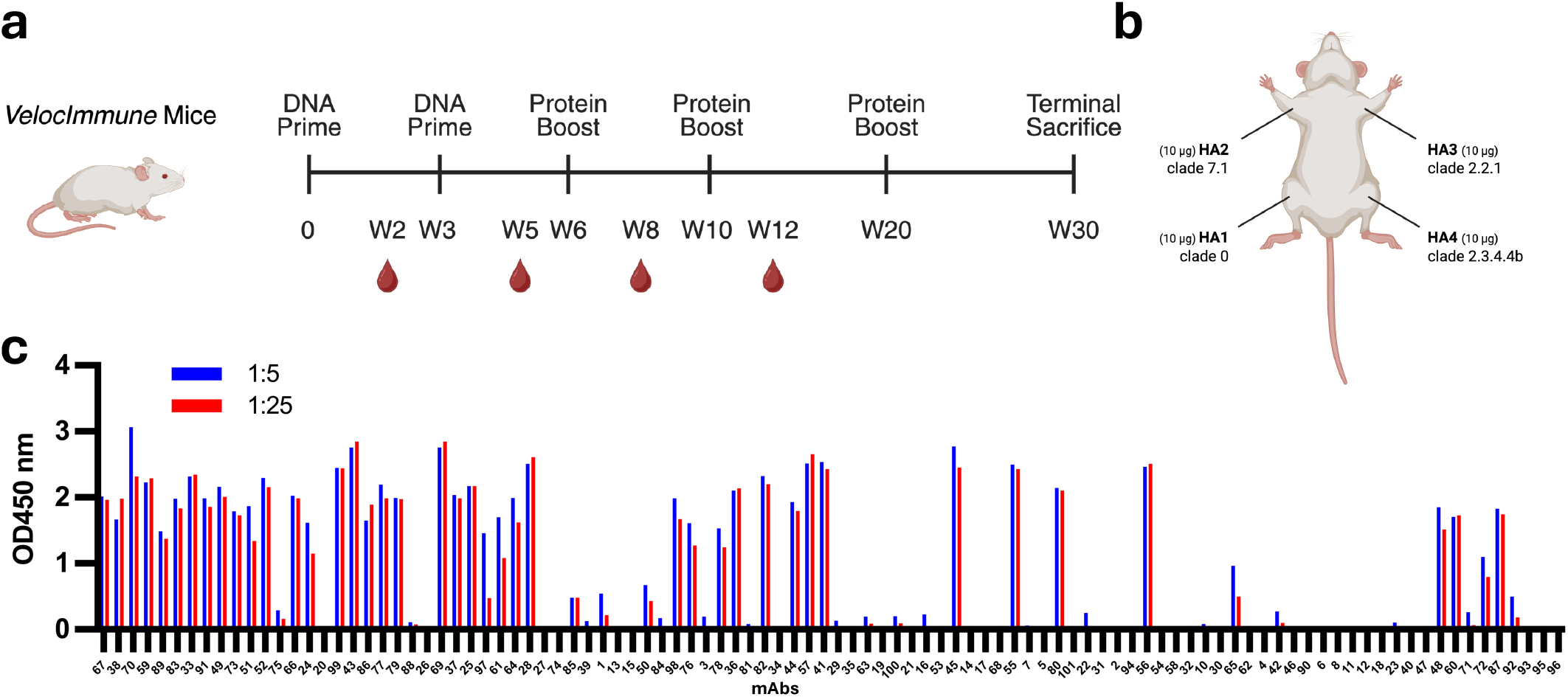
Immunization scheme and supernatant binding screening of selected mAbs. **a.** Immunization schedule and H5 antigens used to immunize VelocImmune® mice. **b.** Schematic of the multi-limb immunization strategy using four selected H5 antigens. **c.** Supernatant binding screen of selected monoclonal antibodies to recombinant A/Texas/37/2024 HA.

**Figure S2.**
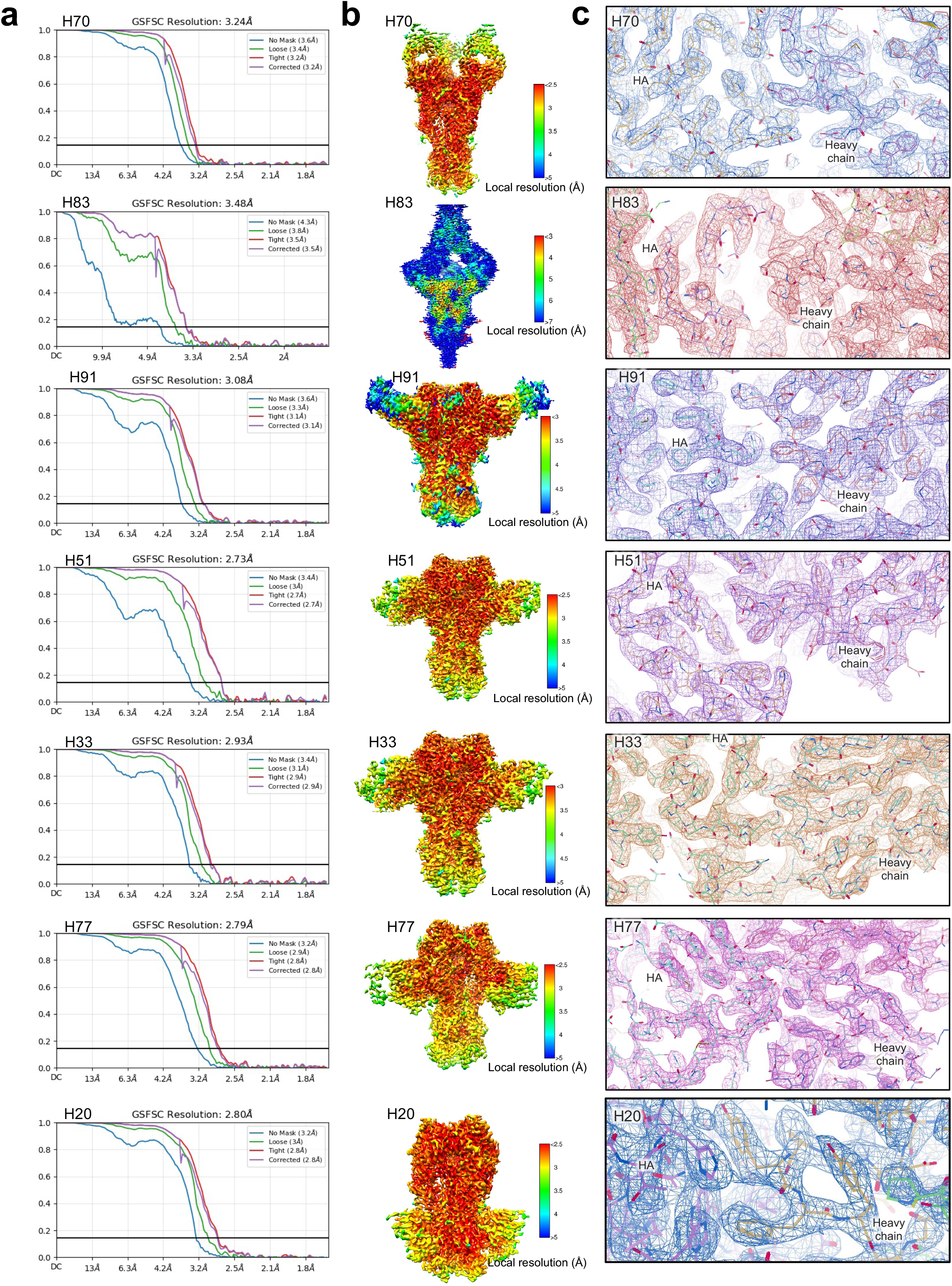
Cryo-EM data for H series Fab:HA complexes. **a**. FSC curves **b**. Side view of 3D volume and local density resolution in Angstroms **c**. Representative density of epitope : paratope interface

**Figure S3.**
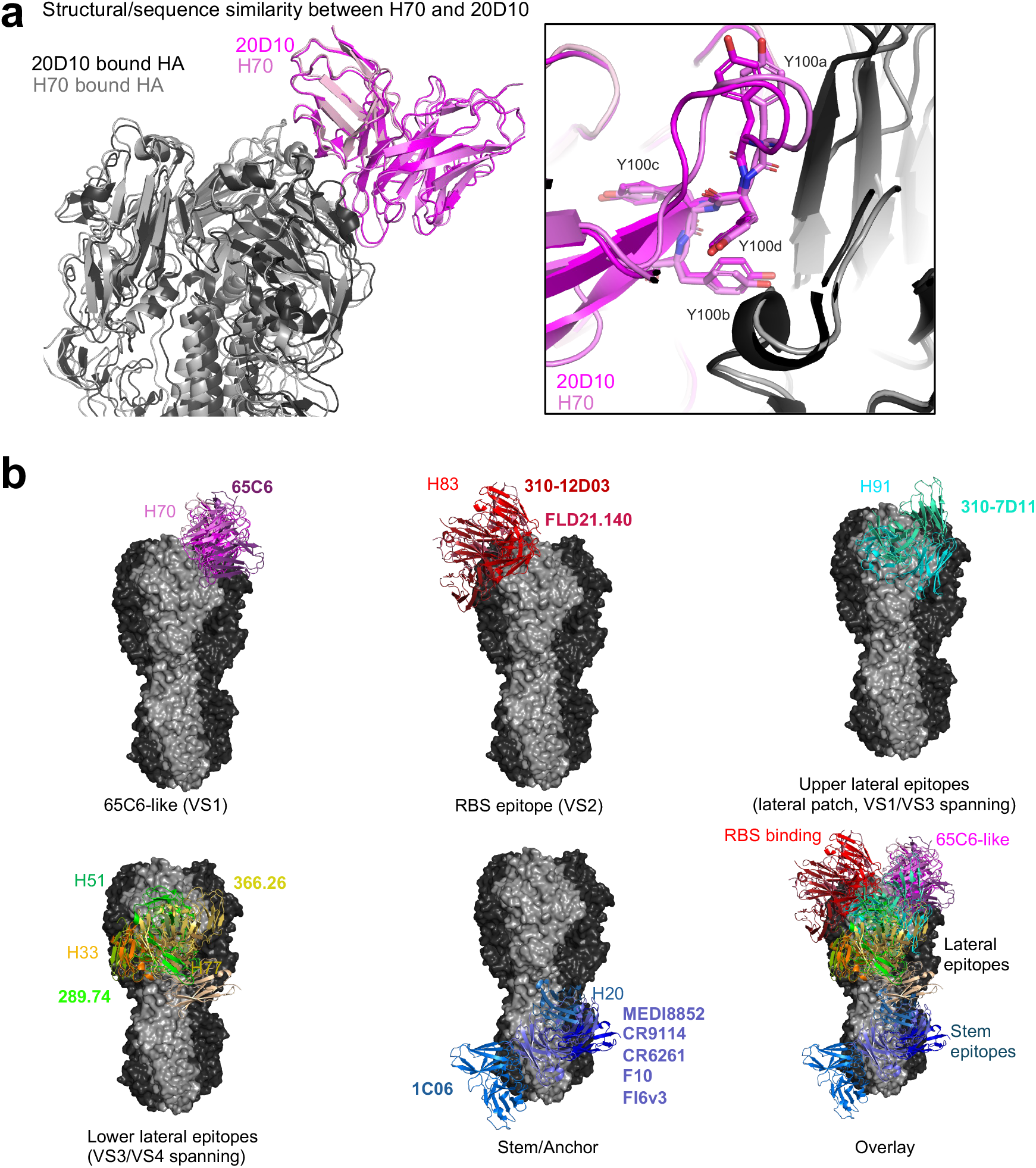
Antibody panel for antigenicity tracking of H5N1 influenza. **a.** Structural/sequence similarity between H70 and 20D10. **b.** Binding sites of a selected antibody panel used to trace H5N1 evolution. Previously published mAbs are shown in bold. CR9114, CR6261, F10, and FI6v3 are not shown but bind to the same epitope as MEDI8852.

**Figure S4.**
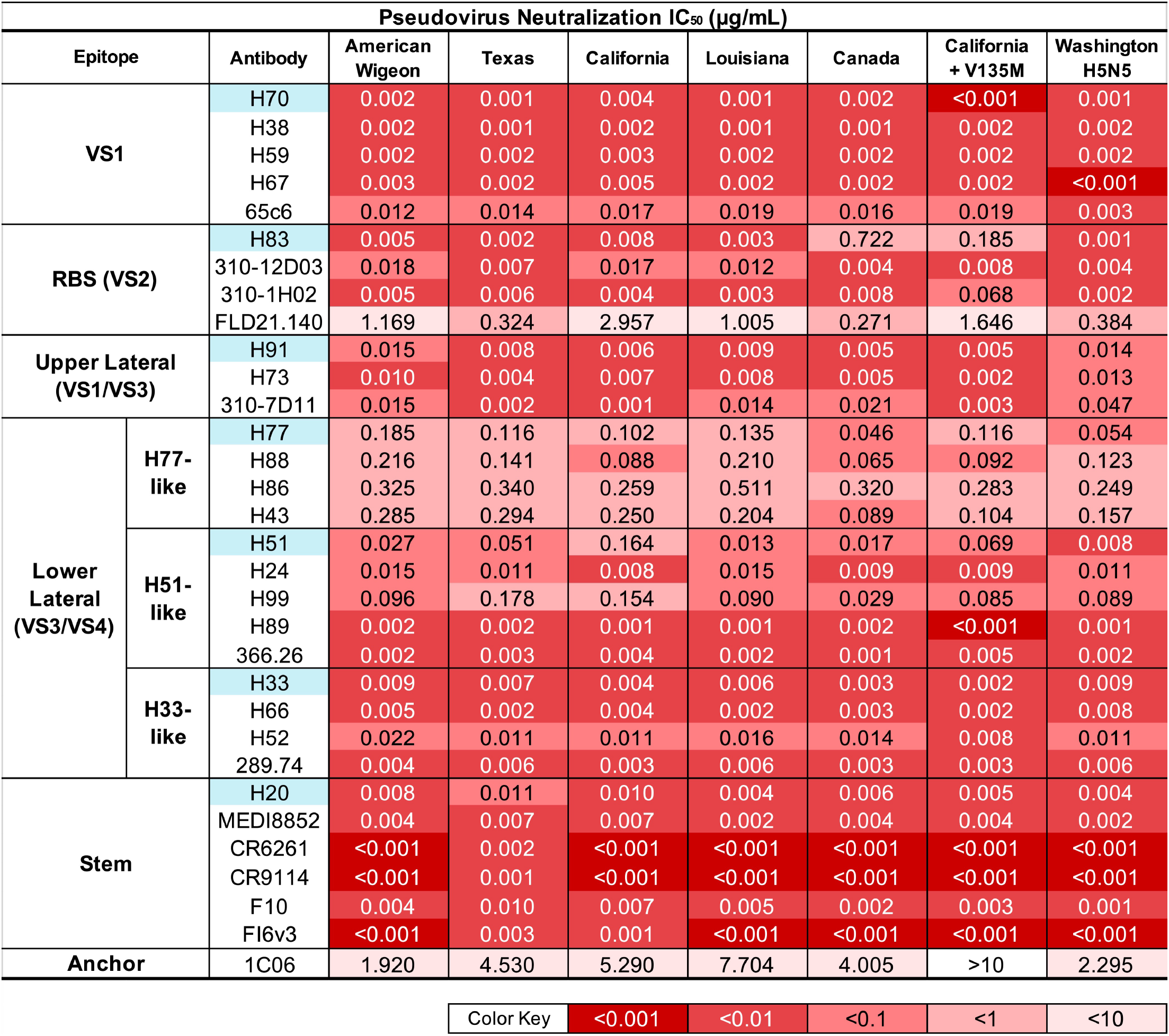
IC50 of neutralizing antibody panel against 2.3.4.4b variants. American Wigeon refers to A/American_Wigeon/South_Carolina/AH0195145/2021; Texas as A/Texas/37/2024; California as A/California/173/2024; Louisiana as A/Louisiana/12/2024; Canada as A/British_Columbia/PHL-2032/2024; California + V135M as A/California/148/2024; Washington H5N5 as A/Washington/2148/2025.

**Figure S5.**
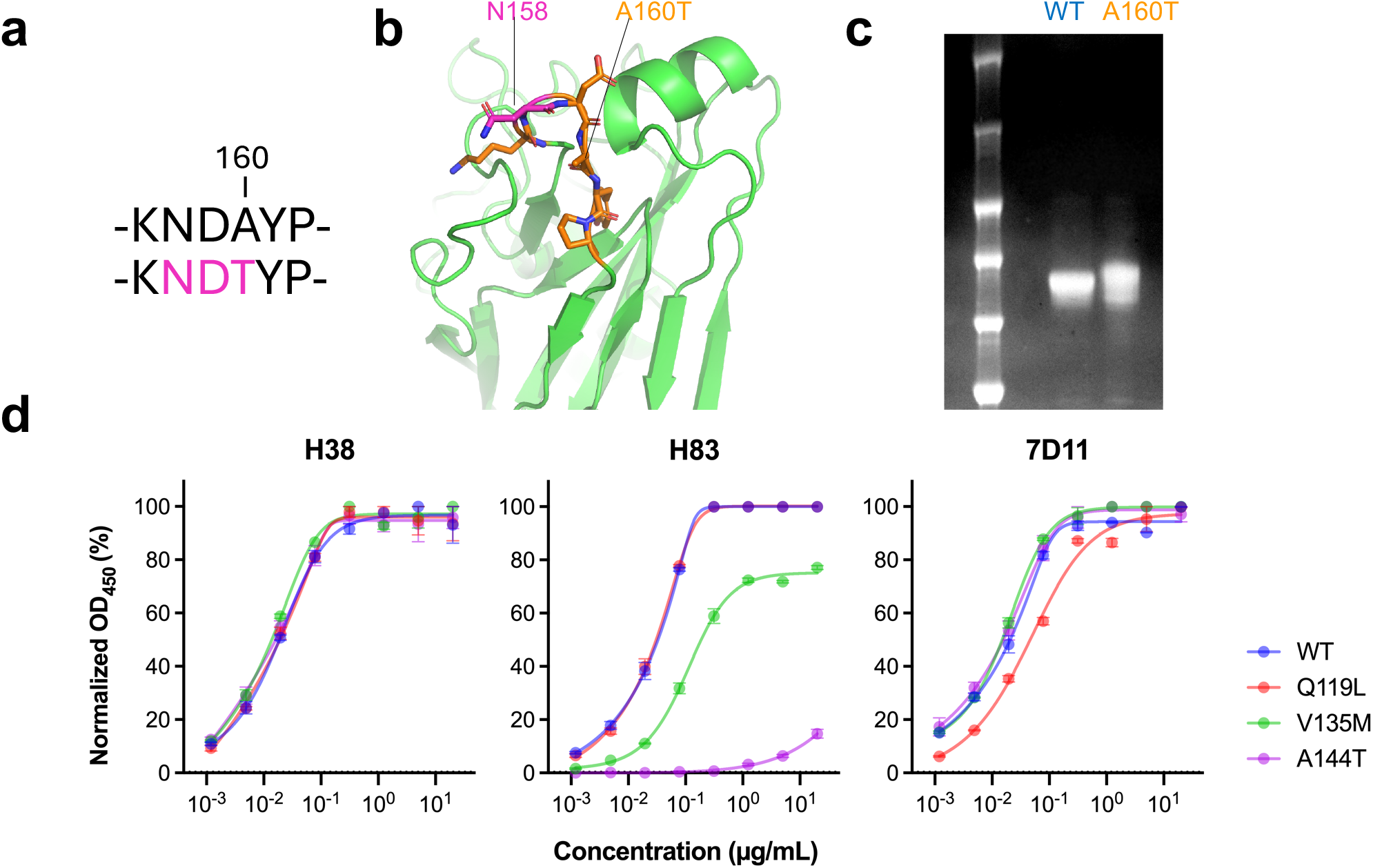
Experimental validation of candidate escape substitutions in A/Texas/37/2024 HA. **a.** HA residues 157-162 in wild-type (-KNDAYP-) and A160T (-KNDTYP-) HA, illustrating the generation of an N-linked glycosylation sequon at Asn158 (magenta). Residues are numbered according to H3 numbering. **b.** Structural locations of Asn158 (magenta) and Ala160 (orange) on a single protomer of the A/Texas/37/2024 HA trimer. **c.** SDS-PAGE analysis of purified wild-type (A160) and mutant (T160) HA proteins. **d.** ELISA binding of H38, H83, and 310-7D11 to wild-type A/Texas/37/2024 HA and HA proteins carrying individual Q119L, V135M, or A144T substitutions. OD450 values were normalized to the maximum signal for each antibody. Data are presented as mean ± SEM of technical duplicates.

## References

1. Li, K.S., Guan, Y., Wang, J., Smith, G.J.D., Xu, K.M., Duan, L., Rahardjo, A.P., Puthavathana, P., Buranathai, C., Nguyen, T.D., et al. (2004). Genesis of a highly pathogenic and potentially pandemic H5N1 influenza virus in eastern Asia. Nature 430, 209–213. 10.1038/nature02746.

2. Xie, R., Edwards, K.M., Wille, M., Wei, X., Wong, S.-S., Zanin, M., El-Shesheny, R., Ducatez, M., Poon, L.L.M., Kayali, G., et al. (2023). The episodic resurgence of highly pathogenic avian influenza H5 virus. Nature 622, 810–817. 10.1038/s41586-023-06631-2.

3. Yuen, K.Y., Chan, P.K.S., Peiris, M., Tsang, D.N.C., Que, T.L., Shortridge, K.F., Cheung, P.T., To, W.K., Ho, E.T.F., Sung, R., et al. (1998). Clinical features and rapid viral diagnosis of human disease associated with avian influenza A H5N1 virus. The Lancet 351, 467–471. 10.1016/S0140-6736(98)01182-9.

4. Claas, E.C., Osterhaus, A.D., Beek, R. van, Jong, J.C.D., Rimmelzwaan, G.F., Senne, D.A., Krauss, S., Shortridge, K.F., and Webster, R.G. (1998). Human influenza A H5N1 virus related to a highly pathogenic avian influenza virus. The Lancet 351, 472–477. 10.1016/S0140-6736(97)11212-0.

5. Garg, S., Reinhart, K., Couture, A., Kniss, K., Davis, C.T., Kirby, M.K., Murray, E.L., Zhu, S., Kraushaar, V., Wadford, D.A., et al. (2025). Highly Pathogenic Avian Influenza A(H5N1) Virus Infections in Humans. N. Engl. J. Med. 392, 843–854. 10.1056/NEJMoa2414610.

6. Damodaran, L., Jaeger, A.S., and Moncla, L.H. (2026). Ecology and spread of the North American H5N1 epizootic. Nature 649, 432–441. 10.1038/s41586-025-09737-x.

7. Krammer, F., Hermann, E., and Rasmussen, A.L. (2025). Highly pathogenic avian influenza H5N1: history, current situation, and outlook. J. Virol. 99, e02209–24. 10.1128/jvi.02209-24.

8. Caserta, L.C., Frye, E.A., Butt, S.L., Laverack, M., Nooruzzaman, M., Covaleda, L.M., Thompson, A.C., Koscielny, M.P., Cronk, B., Johnson, A., et al. (2024). Spillover of highly pathogenic avian influenza H5N1 virus to dairy cattle. Nature 634, 669–676. 10.1038/s41586-024-07849-4.

9. CDC (2026). A(H5) Bird Flu: Current Situation. Avian Influenza Bird Flu. https://www.cdc.gov/bird-flu/situation-summary/index.html.

10. Jassem, A.N., Roberts, A., Tyson, J., Zlosnik, J.E.A., Russell, S.L., Caleta, J.M., Eckbo, E.J., Gao, R., Chestley, T., Grant, J., et al. (2025). Critical Illness in an Adolescent with Influenza A(H5N1) Virus Infection. N. Engl. J. Med. 392, 927–929. 10.1056/NEJMc2415890.

11. Nguyen, T.-Q., Hutter, C.R., Markin, A., Thomas, M., Lantz, K., Killian, M.L., Janzen, G.M., Vijendran, S., Wagle, S., Inderski, B., et al. (2025). Emergence and interstate spread of highly pathogenic avian influenza A(H5N1) in dairy cattle in the United States. Science 388, eadq0900. 10.1126/science.adq0900.

12. Crespo-Bellido, A., Trovão, N.S., Maksiaev, A., Baele, G., Dellicour, S., and Nelson, M.I. (2025). Emergence of D1.1 reassortant H5N1 avian influenza viruses in North America. Preprint at bioRxiv, 10.64898/2025.12.19.695329 https://doi.org/10.64898/2025.12.19.695329.

13. Harrington, W.N., Signore, A., Kercher, L., Giacinti, J.A., Kandeil, A., Ahlstrom, C.A., Bevins, S., Crossley, B., Eure, K., Fabrizio, T.P., et al. (2026). Rapid expansion of genotype D1.1 A(H5N1) influenza viruses in wild birds across North America during the 2024 migratory season. Nat. Med. 32, 1661–1665. 10.1038/s41591-026-04300-1.

14. Lite, T.-L., Goya, S., Davis, M.L., Morris, A.E., Bryson-Cahn, C., Vengerovsky, A., Corpuz, C.G., Nunley, B.E., Maenza, J.R., Hawn, T.R., et al. (2026). Human Infection with Highly Pathogenic Avian Influenza A(H5N5) Virus. N. Engl. J. Med. 394, 2274–2276. 10.1056/NEJMc2518178.

15. CDC (2025). Genetic Sequences of Highly Pathogenic Avian Influenza A(H5N1) Viruses Identified in a Person in Louisiana. Avian Influenza Bird Flu. https://www.cdc.gov/bird-flu/spotlights/h5n1-response-12232024.html.

16. Avian Influenza A(H5N1) - Mexico https://www.who.int/emergencies/disease-outbreak-news/item/2025-DON564.

17. Mostafa, A., Nogales, A., and Martinez-Sobrido, L. (2025). Highly pathogenic avian influenza H5N1 in the United States: recent incursions and spillover to cattle. Npj Viruses 3, 54. 10.1038/s44298-025-00138-5.

18. Nair, M.S., Hong, H., Chong, S., Huang, Y., and Ho, D.D. (2026). Serum neutralisation of H5N1 clade 2.3.4.4b influenza is largely mediated by neuraminidase-directed antibodies. Lancet Microbe 7. 10.1016/j.lanmic.2026.101431.

19. Dadonaite, B., Ahn, J.J., Ort, J.T., Yu, J., Furey, C., Dosey, A., Hannon, W.W., Baker, A.L.V., Webby, R.J., King, N.P., et al. (2024). Deep mutational scanning of H5 hemagglutinin to inform influenza virus surveillance. PLOS Biol. 22, e3002916. 10.1371/journal.pbio.3002916.

20. Belser, J.A. (2025). A pandemic toolbox for clade 2.3.4.4b A(H5N1) influenza virus risk assessment. Lancet Microbe 0. 10.1016/j.lanmic.2025.101240.

21. Ford, C.T., Yasa, S., Obeid, K., Jaimes, R., Tomezsko, P.J., Guirales-Medrano, S., White, R.A., and Janies, D. (2025). Large-scale computational modelling of H5 influenza variants against HA1-neutralising antibodies. eBioMedicine 114. 10.1016/j.ebiom.2025.105632.

22. Mellis, I.A., Wu, M., Hong, H., Tzang, C.-C., Bowen, A., Wang, Q., Gherasim, C., Pierce, V.M., Shah, J.G., Purpura, L.J., et al. (2025). Antibody evasion and receptor binding of SARS-CoV-2 LP.8.1.1, NB.1.8.1, XFG, and related subvariants. Cell Rep. 44. 10.1016/j.celrep.2025.116440.

23. Hong, H., Wu, M., Chong, S., Yu, J., Mellis, I.A., Huang, C.-Y., Wang, Q., Huang, Y., Guo, Y., and Ho, D.D. (2026). A Monoclonal Antibody Panel to Track Antigenic Evolution of SARS-CoV-2 JN.1 Subvariants. Preprint at bioRxiv, 10.64898/2026.05.29.728770 https://doi.org/10.64898/2026.05.29.728770.

24. Zuo, T., Sun, J., Wang, G., Jiang, L., Zuo, Y., Li, D., Shi, X., Liu, X., Fan, S., Ren, H., et al. (2015). Comprehensive analysis of antibody recognition in convalescent humans from highly pathogenic avian influenza H5N1 infection. Nat. Commun. 6, 8855. 10.1038/ncomms9855.

25. Abu-Shmais, A.A., Freeman, G., Creanga, A., Vukovich, M.J., Malla, T., Mantus, G.E., Shimberg, G.D., Gillespie, R.A., Guerra Canedo, V., Dadonaite, B., et al. (2025). Cross-neutralizing and potent human monoclonal antibodies against historical and emerging H5Nx influenza viruses. Nat. Microbiol. 10, 2903–2918. 10.1038/s41564-025-02137-x.

26. Alzua, G.P., León, A.N., Yellin, T., Bhavsar, D., Loganathan, M., Bushfield, K., Brouwer, P.J.M., Rodriguez, A.J., Jeevan, T., Webby, R., et al. (2025). Human monoclonal antibodies that target clade 2.3.4.4b H5N1 hemagglutinin. Nat. Commun. 10.1038/s41467-025-66829-y.

27. Bangaru, S., Lang, S., Schotsaert, M., Vanderven, H.A., Zhu, X., Kose, N., Bombardi, R., Finn, J.A., Kent, S.J., Gilchuk, P., et al. (2019). A Site of Vulnerability on the Influenza Virus Hemagglutinin Head Domain Trimer Interface. Cell 177, 1136–1152.e18. 10.1016/j.cell.2019.04.011.

28. Hu, H., Voss, J., Zhang, G., Buchy, P., Zuo, T., Wang, L., Wang, F., Zhou, F., Wang, G., Tsai, C., et al. (2012). A Human Antibody Recognizing a Conserved Epitope of H5 Hemagglutinin Broadly Neutralizes Highly Pathogenic Avian Influenza H5N1 Viruses. J. Virol. 86, 2978– 2989. 10.1128/jvi.06665-11.

29. Kallewaard, N.L., Corti, D., Collins, P.J., Neu, U., McAuliffe, J.M., Benjamin, E., Wachter-Rosati, L., Palmer-Hill, F.J., Yuan, A.Q., Walker, P.A., et al. (2016). Structure and Function Analysis of an Antibody Recognizing All Influenza A Subtypes. Cell 166, 596–608. 10.1016/j.cell.2016.05.073.

30. Feldman, J., Ramos, A.S.F., Vu, M., Maurer, D.P., Rosado, V.C., Lingwood, D., Bajic, G., and Schmidt, A.G. (2025). Human naïve B cells recognize prepandemic influenza virus hemagglutinins. Sci. Immunol. 10, eado9572. 10.1126/sciimmunol.ado9572.

31. Murphy, A.J., Macdonald, L.E., Stevens, S., Karow, M., Dore, A.T., Pobursky, K., Huang, T.T., Poueymirou, W.T., Esau, L., Meola, M., et al. (2014). Mice with megabase humanization of their immunoglobulin genes generate antibodies as efficiently as normal mice. Proc. Natl. Acad. Sci. 111, 5153–5158. 10.1073/pnas.1324022111.

32. Wang, Q., Guo, Y., Casner, R.G., Yu, J., Nair, M.S., Ho, J., Reddem, E.R., Mellis, I.A., Wu, M., Tzang, C.-C., et al. (2025). Optimizing a human monoclonal antibody for better neutralization of SARS-CoV-2. Nat. Commun. 16, 6195. 10.1038/s41467-025-61472-z.

33. Liu, L., Wang, P., Nair, M.S., Yu, J., Rapp, M., Wang, Q., Luo, Y., Chan, J.F.-W., Sahi, V., Figueroa, A., et al. (2020). Potent neutralizing antibodies against multiple epitopes on SARS-CoV-2 spike. Nature 584, 450–456. 10.1038/s41586-020-2571-7.

34. Liu, L., Casner, R.G., Guo, Y., Wang, Q., Iketani, S., Chan, J.F.-W., Yu, J., Dadonaite, B., Nair, M.S., Mohri, H., et al. (2023). Antibodies targeting a quaternary site on SARS-CoV-2 spike glycoprotein prevent viral receptor engagement by conformational locking. Immunity 56, 2442–2455.e8. 10.1016/j.immuni.2023.09.003.

35. Morano, N.C., Guo, Y., Becker, J.E., Li, Z., Yu, J., Ho, D.D., Shapiro, L., and Kwong, P.D. (2025). Structure of a zoonotic H5N1 hemagglutinin reveals a receptor-binding site occupied by an auto-glycan. Structure 33, 228–233.e3. 10.1016/j.str.2025.01.001.

36. Dunbar, J., Krawczyk, K., Leem, J., Marks, C., Nowak, J., Regep, C., Georges, G., Kelm, S., Popovic, B., and Deane, C.M. (2016). SAbPred: a structure-based antibody prediction server. Nucleic Acids Res. 44, W474–W478. 10.1093/nar/gkw361.

37. Emsley, P., Lohkamp, B., Scott, W.G., and Cowtan, K. (2010). Features and development of *Coot*. Acta Crystallogr. D Biol. Crystallogr. 66, 486–501. 10.1107/S0907444910007493.

